# Decitabine Reactivation of FoxM1-Dependent Endothelial Regeneration and Vascular Repair for Potential Treatment of Elderly ARDS and COVID-19 Patients

**DOI:** 10.1101/2021.04.29.442061

**Authors:** Xiaojia Huang, Xianming Zhang, Narsa Machireddy, Gökhan M. Mutlu, Yun Fang, David Wu, You-Yang Zhao

## Abstract

Aging is a major risk factor of high incidence and increased mortality of acute respiratory distress syndrome (ARDS) and COVID-19. We repot that aging impairs the intrinsic FoxM1-dependent endothelial regeneration and vascular repair program and causes persistent lung injury and high mortality following sepsis. Therapeutic gene transduction of *FOXM1* in vascular endothelium or treatment with FDA-approved drug Decitabine was sufficient to reactivate FoxM1-dependent lung endothelial regeneration in aged mice, reverse aging-impaired resolution of inflammatory injury, and promote survival. In COVID-19 lung autopsy samples, FOXM1 expression was not induced in vascular endothelial cells of elderly patients in contrast to mid-age patients. Thus, Decitabine reactivation of FoxM1-dependent vascular repair represents a potential effective therapy for elderly COVID-19 and non-COVID-19 ARDS patients.

## INTRODUCTION

The global COVID-19 pandemic has signified the urgency and importance of understanding the molecular and cellular mechanisms of endothelial regeneration and vascular repair in the pathogenesis of acute respiratory distress syndrome (ARDS). ARDS is a form of acute-onset hypoxemic respiratory failure with bilateral pulmonary infiltrates, which is caused by acute inflammatory edema of the lungs not attributable to left heart failure (*1–3*). The common causes of ARDS include sepsis, pneumonia, inhalation of harmful substance, burn, major trauma with shock and massive transfusion. Sepsis with annual U.S. incidence of over 750,000 is the most common cause. In the current COVID-19 pandemic, COVID-19 has also found to be the major cause of ARDS. Endothelial injury characterized by persistently increased lung microvascular permeability resulting in protein-rich lung edema is a hallmark of acute lung injury (ALI)/ARDS (*4–7*). Despite recent advances on the understanding of the pathogenesis, there are currently no effective pharmacological or cell-based treatment of the disease and the mortality remains as high as 40% (*1–3*). Compared to young adult patients, the incidence of ARDS resulting from sepsis and pneumonia in elderly patients (≥ 65 yr) is as much as 19-fold greater and the mortality is up to 10-fold greater (*1, 8–11*). Similarly, the hospitalization and mortality rates of elderly COVID-19 patients are much higher than young adults (*12–15*). In one study, it has been shown that 43% of COVID-19 patients aged ≥ 70 years died compared to 8.7% of 40-59 years old and 2.8% of 20-39 years old patients died (*14*). Severe vascular endothelial injury derived from direct viral effects and perivascular inflammation is a characteristic feature of COVID-19 lung injury (*16–19*) and COVID-19 ARDS is considered as a vascular endotype of ARDS (*20–22*). However, little is known about how aging influences mechanisms of endothelial regeneration and resulting restoration of vascular homeostasis. The underlying causes of aging-related high incidence and mortality of ARDS and COVID-19 are poorly understood and thus there is no effective therapy to prevent and treat COVID-19 and non-COVID-19 ARDS.

Here we sought to define the cell source of origin mediating lung endothelial regeneration following sepsis injury by genetic lineage tracing, determine how aging affects this process and resolution of inflammatory lung injury, and delineate the underlying molecular mechanisms. Our studies demonstrate that aging impairs the intrinsic endothelial regeneration and vascular repair program and thus resolution of inflammation. Forced expression of the forkhead transcription factor FoxM1 (*23–26*) in lung ECs (transgenic or non-viral delivery of plasmid DNA) in aged mice is sufficient to re-activate lung endothelial regeneration and vascular repair and promote survival following sepsis. FoxM1 expression was induced in pulmonary vascular ECs of mid-aged COVID-19 patients but not in elderly patients. Importantly, repurposing an FDA-approved drug Decitabine could reactivate FoxM1-dependent endothelial regeneration and promote survival in aged mice. Thus, our studies demonstrate repurposing Decitabine to activate FoxM1-dependent endothelial regeneration and vascular repair represents a potential novel and effective therapy of ARDS and severe COVID-19 in elderly patients.

## RESULTS

### Aging impairs intrinsic lung endothelial regeneration following polymicrobial sepsis-induced injury

The major pathogenic feature of ALI/ARDS leading to deterioration of vascular barrier function is the precipitous loss of ECs (*27–30*). To trace the changes of pulmonary ECs following sepsis challenge, we performed lineage tracing on pulmonary ECs using a tamoxifen-inducible mTmG/*EndoSCL*-Cre^ERT2^ mouse (**fig. S1A**). 95% of lung ECs (CD45^−^CD31^+^) were labeled with green fluorescent protein (GFP) after tamoxifen treatment (**fig. S1, B** **and** **C**) whereas <5% of GFP^+^ cells were either CD45^+^ cells (leukocytes) or CD31^−^ cells (non-ECs) (**fig. S1, D** **to** **G**).

Fluorescence imaging revealed GFP^+^ ECs in capillaries and along the inner surfaces of blood vessels but not bronchioles (**Fig. 1A**). In young adult mice (3-5 mos. old), at 48h post-cecal ligation and puncture (CLP), which causes lethal peritonitis and polymicrobial sepsis, a well-recognized clinically relevant murine model of sepsis (*31, 32*), the presence of GFP^+^ ECs was noticeably disrupted along the blood vessel inner surfaces, consistent with loss of ECs seen in patients and animal models; by 144h post-CLP, the blood vessel inner wall was nicely lined with GFP^+^ ECs again (**fig. S2**). To quantify the changes of pulmonary EC numbers over the course of sepsis-induced injury and recovery, we measured the percentage of CD45^−^GFP^+^ cells in the whole lung by flow cytometry analysis (FACS) in young adult mice. In sham animals, ~40% of pulmonary CD45^−^ cells were GFP^+^. At 48h post-CLP, this number had dropped to 25%, but was followed by a steady return to baseline levels by 144h (**Fig. 1, B** and **C**). However, the CD45^+^GFP^+^ cell population was remained at steady minimal levels at various times (**fig. S3**), indicating CD45^+^GFP^+^ cells were not involved in endothelial regeneration. Bone marrow cell transplantation study further demonstrated that bone marrow-derived cells were not attributable to endothelial regeneration as the transplanted GFP^+^ population remained steady (**fig. S4**). Together, these data demonstrate that lung resident ECs are the cell source for endothelial regeneration in young adult mice following sepsis-induced injury.

**Fig. 1.**
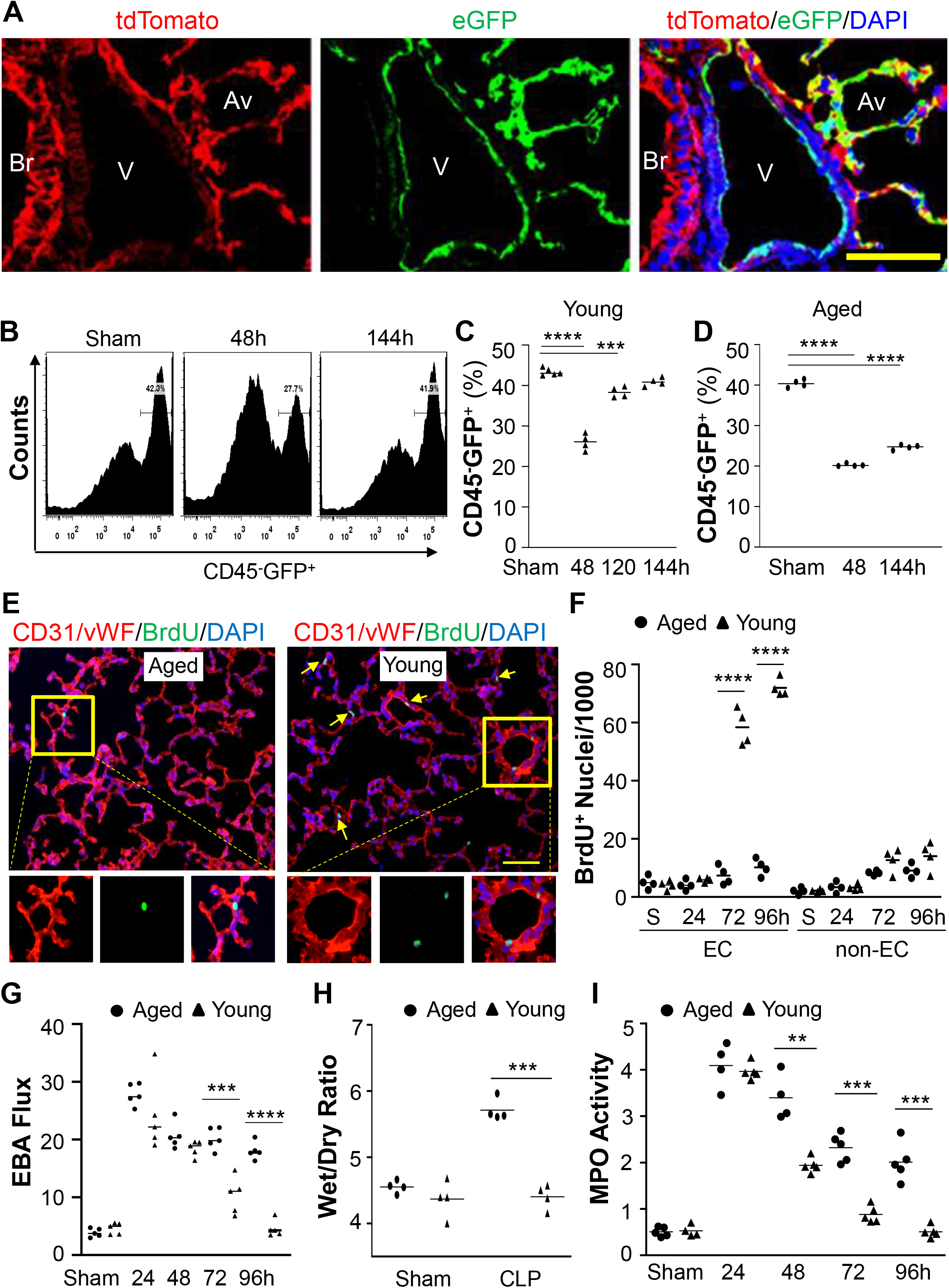
Defective endothelial regeneration and vascular repair in aged lungs following polymicrobial sepsis. (**A**) Representative confocal images of lungs of young adult mice showing GFP-labeling of vascular ECs. V=vessel, Br=bronchiole, Av-alveolus. Scale bar, 20 μm. (**B**, **C**) FACS analysis demonstrating loss of lung GFP^+^ ECs at 48h post-CLP and steady recovery of GFP^+^ ECs during the repair phase in young adult mice. Lung cells were CD45^−^ gated, and GFP^+^ population were quantified. (**D**) FACS analysis showing impaired recovery of lung GFP^+^ cells following CLP in aged mice (19-21 mos. old). (**E**) Representative micrographs of BrdU immunostaining showing defective EC proliferation in aged lungs. Cryosections of lungs collected at 96h post-CLP were immunostained with anti-BrdU antibody to identify proliferating cells (green) and with anti-CD31 and vWF antibodies to identify ECs (red). Arrows point to proliferating ECs. Scale bar, 50 μm. (**F**) Quantification of cell proliferation in mouse lungs. (**G**) Lung vascular permeability assessed by EBA extravasation assay. (**H**) Lung wet/dry weight ratio. At 96h post-CLP, lung tissues were collected and dried at 60°C for 3 days for calculation of wet/dry ratio. (**I**) MPO activities in lung tissues. Bars represent means. \*\**P*<0.01; \*\*\**P*<0.001, \*\*\*\**P*<0.0001. Unpaired *t* test (parametric).

FACS analysis revealed that the lung GFP^+^ EC population was markedly decreased at 48h post-CLP in aged (19-21 mos.) mice as observed in young adult mice (**Fig. 1D**). However, in contrast to young adult mice (**Figure 1C**), the GFP^+^ EC population in aged mice failed to recover and remained low at 144h post-CLP (**Fig. 1D**). Thus, aging impaired the intrinsic endothelial regeneration program following sepsis challenge.

### Impaired endothelial regeneration leads to persistent inflammatory lung injury in aged mice following polymicrobial sepsis

We next employed bromodeoxyuridine (BrdU) pulse assay to assess cell proliferation. Anti-BrdU immunostaining revealed defective lung endothelial proliferation in aged mice in contrast to young adult mice during the recovery phase (e.g., 72 and 96h post-CLP) (**Fig. 1, E** and **F**). Accordingly, Evans blue-conjugated albumin (EBA) assay, a measurement of vascular permeability to protein showed persistent vascular leak indicating impaired vascular repair in the lungs of aged mice whereas vascular permeability returned to basal levels at 96h post-CLP in young adult mice (**Fig. 1G**). The aged lungs also exhibited marked edema measured by greater lung wet/dry weight ratio at 72h post-CLP (**Fig. 1H**) and impaired resolution of inflammation during the recovery phase evident by persistently elevated lung myeloperoxidase (MPO) activity (**Fig. 1I**), indicative of neutrophil sequestration, and increased expression of proinflammatory cytokines in lung tissue (**fig. S5**).

### Aging impairs lung vascular repair and resolution of inflammation following endotoxemia

To determine if aged mice also exhibit impaired vascular repair following endotoxemia, aged (19-21 mos.) and young (3-5 mos.) mice were challenged with lipopolysaccharide (LPS). Given that aged mice exhibited greater lung injury indicated by greater EBA flux and MPO activity at 24h post-LPS compared to young adult mice in response to the same dose of LPS (data not shown), we challenged the aged mice with a lower dose of LPS (e.g., 1.0 mg/kg) to induce a similar degree of injury during the injury phase (e.g., 24h) as seen in young adult mice with 2.5 mg/kg of LPS (**Fig. 2A**). EBA flux in young adult mice was reduced at 48h and returned to basal levels at 72h post-LPS whereas it remained elevated in aged lungs demonstrating defective vascular repair in aged lungs (**Fig. 2A**). Consistently, aged lungs exhibited edema at 72h post-LPS, which was not observed in young adult mice (**Fig. 2B**). MPO activity remained elevated in aged lungs at 72h post-LPS (**Fig. 2C**). Furthermore, quantitative RT-PCR analysis demonstrated markedly elevated expression of proinflammatory genes including *Tnf* and *Il6* in lungs of aged but not young adult mice at 72h post-LPS (**fig. S6**). Together, these data demonstrated impaired vascular repair and resolution of inflammation in aged lungs following LPS challenge.

**Fig. 2.**
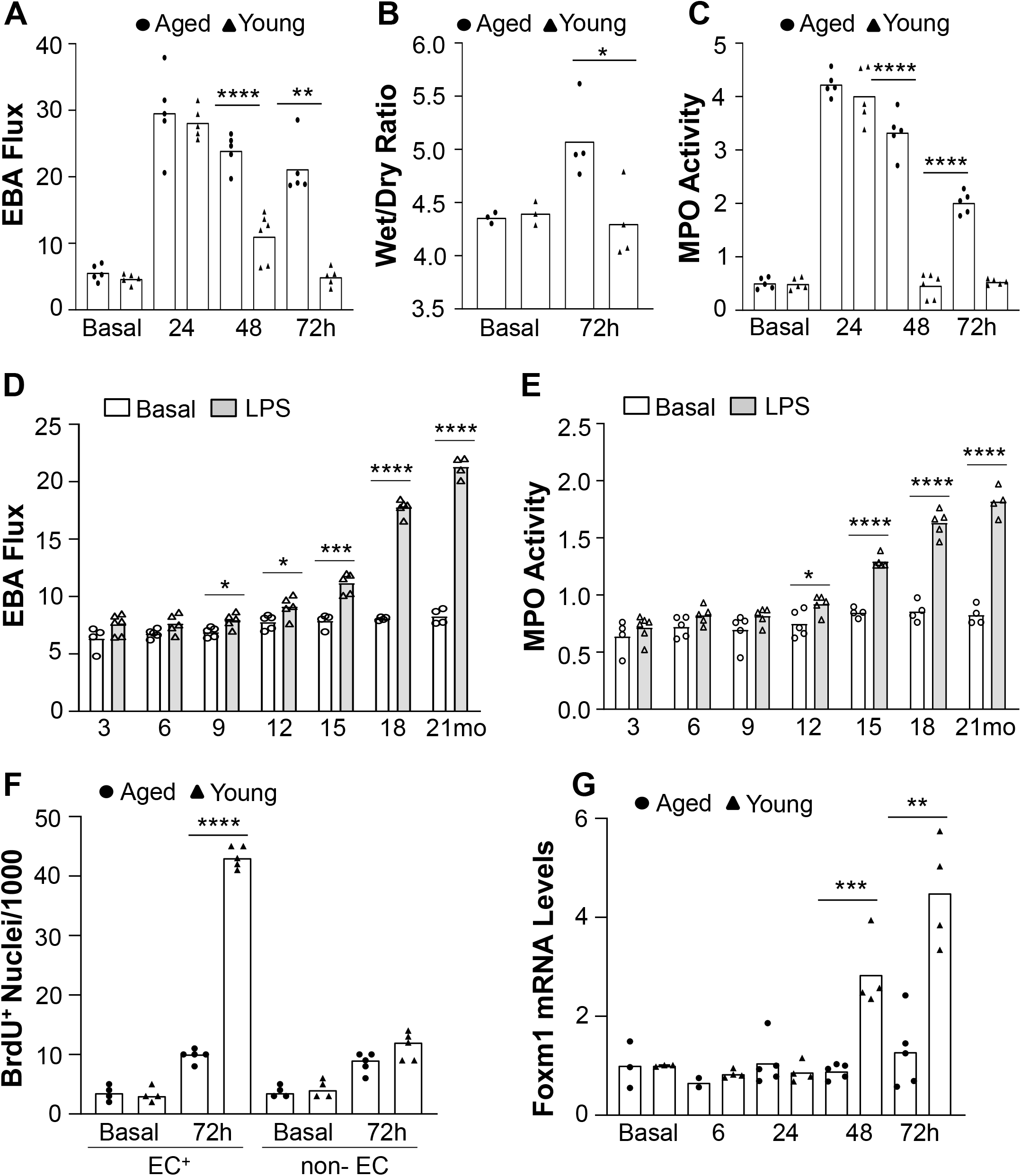
Aging impairment of vascular repair and resolution of inflammatory lung injury following LPS challenge. (**A**) Persistent increase of lung vascular permeability in aged mice following LPS challenge (i.p.). (**B**) Lung edema in aged mice at 72h post-LPS. (**C**) Sustained increase of MPO activity in aged lungs following LPS challenge. (**D**) EBA flux assay demonstrating that aging impaired vascular repair. WT mice at indicated ages were challenged with LPS (mice at age of 3-9 mo. with 2.5 mg/kg, and at age of 12-21 mo. with 1.25 mg/kg LPS). At 72h post-LPS, lungs were collected for EBA extravasation assay. (**E**) Lung MPO activity at 72h post-LPS. (**F**) Quantification of cell proliferation (anti-BrdU^+^ cells) in mouse lungs at basal and 72h post-LPS. (**G**) Quantitative RT-PCR analysis of FoxM1 expression in mouse lungs at indicated times post-LPS. \**P*<0.05; \*\**P*<0.01; \*\*\**P*<0.001; \*\*\*\**P*<0.0001. Unpaired *t* test except Mann-Whitney test for **A** (72h) and **G** (48h).

To further determine how aging affects vascular repair and inflammation resolution, we challenged the mice at various ages (3 to 21 mos. old) with LPS and assessed EBA flux and MPO activity at 72h post-LPS. As shown in **Fig. 2D**, EBA flux in mice at age of 6 mos. or younger returned to basal levels whereas it did not fully recover and maintained at marginally increased levels in 9 and 12 mos. old mice. EBA flux was markedly elevated in 15 mos. old mice and greatly exaggerated in aged mice (18 and 21 mos. old). We also observed similar changes in MPO activity. Lung MPO activity did not return to basal levels at 72h post-LPS in mice starting at age 12 mos. and remained markedly increased in lungs of mice at age 15 mos. or older, indicating impaired resolution of inflammation (**Fig. 2E**). Thus, mice at age 18 mos. or older exhibited severely impaired vascular repair and resolution of inflammation.

### Defective endothelial proliferation and inhibited FoxM1 induction in aged lungs following LPS challenge

Anti-BrdU immunostaining shows a marked increase of endothelial proliferation in the lungs of young adult mice at 72h post-LPS whereas endothelial proliferation in lungs of aged mice was largely inhibited (**Fig. 2F**), indicating impaired endothelial regeneration in aged lungs following LPS challenge. As FoxM1 is a critical reparative transcriptional factor (*27, 33, 34*), we assessed FoxM1 expression in mouse lungs. Foxm1 was markedly induced in the lungs of young adult mice during the recovery phase but not in aged lungs following LPS challenge (**Fig. 2G**). Accordingly, Foxm1 target genes essential for cell cycle progression were not induced in aged lungs (**fig. S7**).

### Normalized vascular repair and inflammation resolution in aged *Foxm1^Tg^* mice

To determine if failure of FoxM1 induction is responsible for the impaired vascular repair and inflammation resolution in aged mice, we employed the *Foxm1^Tg^* mice expressing human *FOXM1* under the control of *Rosa26* promoter (*35*). EBA flux was increased similarly at 24h post-LPS challenge in aged *Foxm1^Tg^* mice compared to aged WT mice, demonstrating similar degree of lung vascular injury (**Fig. 3A**). EBA flux was then reduced at 48h and returned to a level close to basal level at 72h post-LPS in aged *Foxm1^Tg^* mice whereas it was persistently elevated in aged WT mice (**Fig. 3A**). MPO activity was also similarly increased during the injury phase in aged WT and *Foxm1^Tg^* mice and returned to basal levels at 72h post-LPS in aged *Foxm1^Tg^* mice but not in aged WT mice (**Fig. 3B**). These data demonstrate normalized vascular repair and resolution of inflammation in aged *Foxm1^Tg^* mice following LPS challenge. Accordingly, transgenic expression of FoxM1 promoted survival. 70% of aged *Foxm1^Tg^* mice survived in 7 days following LPS challenge whereas all the aged WT mice died (**Fig. 3C**).

**Fig. 3.**
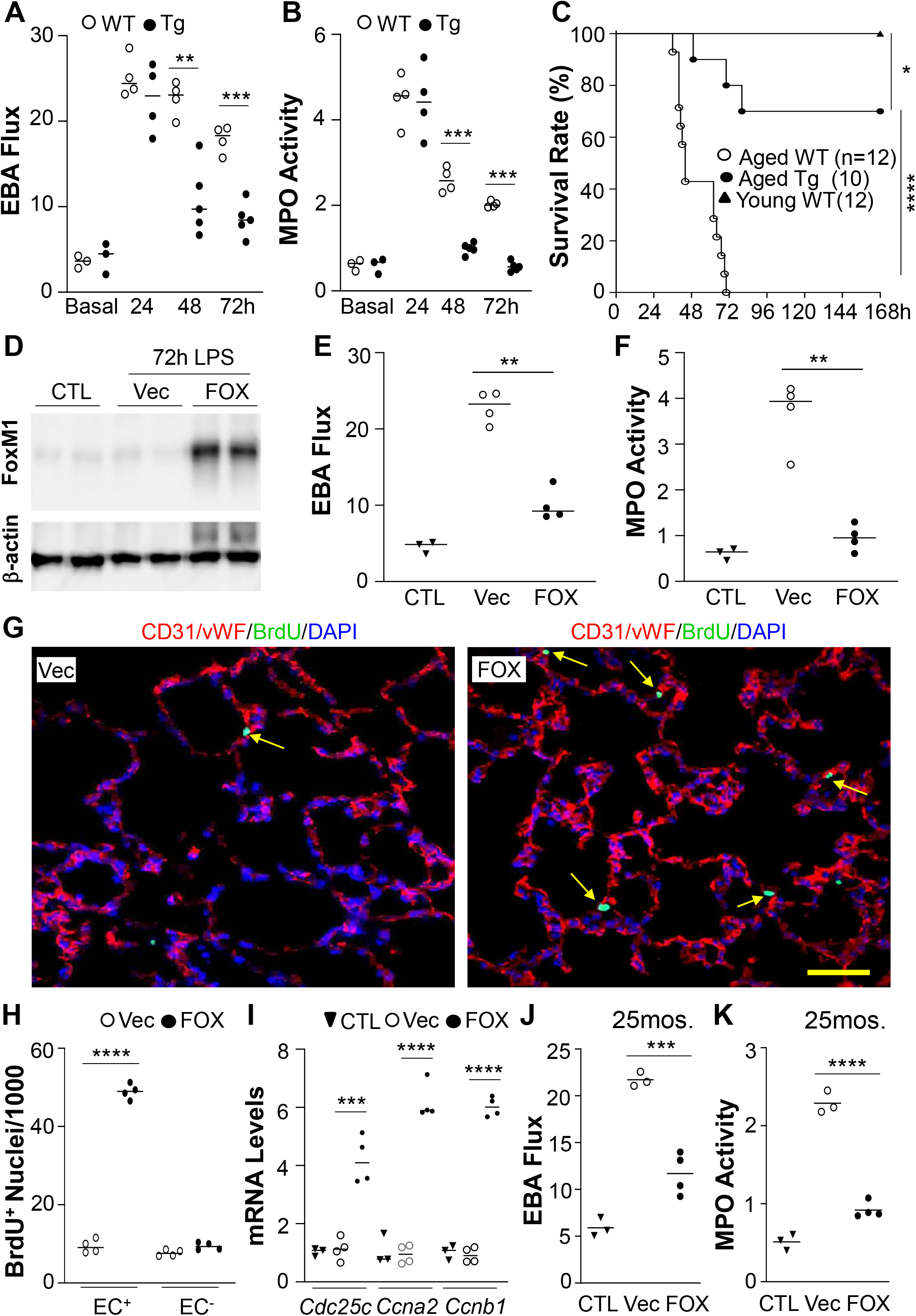
Forced expression of FoxM1 normalized resolution of inflammatory lung injury and promoted survival of aged mice. FoxM1 expression was increased in aged mice by transgene (**A-C**), liposome (**D-I**) or nanoparticle (**J**, **K**):plasmid DNA transduction. Aged WT or *FoxM1^Tg^* mice (21 mos.) were challenged with LPS (i.p.). (**A**) Normalized vascular repair in aged *FOXM1^Tg^* mice. (**B**) Lung MPO activity assessment demonstrating normal resolution of inflammation in aged *FOXM1^Tg^* mice. (**C**) Transgenic expression of FoxM1 promoted survival of aged *FOXM1^Tg^* mice. (**D**) Representative Western blotting demonstrating marked increase of FoxM1 expression in lungs of aged WT mice transduced with *FOXM1* plasmid DNA (FOX) compared to empty vector DNA (Vec). Mixture of liposome: *FOXM1* plasmid DNA (*CDH5* promoter) or vector DNA were administered retro-orbitally to aged WT mice at 12h post-LPS (50μg DNA/mouse). Lungs were collected at 72h post-LPS (**D**-**H**). (**E**) Marked decrease of lung vascular permeability in *FOXM1*-transduced mice. (**F**) Lung MPO activity. (**G, H**) Transient FoxM1 expression in lung ECs of aged WT mice reactivated lung EC proliferation. Arrows point to proliferating ECs. Scale bar, 60μm. (**I**) Quantitative RT-PCR analysis of FoxM1 target genes. (**J, K**) Nanoparticle delivery of *FOXM1* plasmid DNA to elderly mice activated vascular repair and inflammation resolution. Mixture of nanoparticle:DNA was administered retro-orbitally (15μg DNA/mouse) at 24h post-LPS (0.25mg/kg, i.p.). At 96h post-LPS, lung tissues were collected for EBA extravasation assay (**J**) and MPO activity determination (**K**). \**P*<0.05; \*\**P*<0.01; \*\*\**P*<0.001; \*\*\*\**P*<0.0001. Unpaired *t* test (**A**, **B**, **E**-**K**). Log-rank (Mantel-Cox) test (**C**).

### Restored endothelial regeneration and resolution of inflammatory lung injury in aged WT mice by FoxM1 gene transduction

Next, we employed a gene therapy approach to determine if forced FoxM1 expression in lung vascular ECs of aged WT mice after sepsis can reactivate endothelial regeneration and thus restore the defective resolution of inflammatory lung injury. A mixture of liposome:plasmid DNA (*36*) expressing human *FOXM1* under the control of human *CDH5* promoter (EC-specific) or empty vector DNA was administered retro-orbitally to 19-20 mos. old WT mice at 12h post-LPS (established lung injury). At 72h post-LPS, liposome transduction of *FOXM1* plasmid DNA resulted in a marked increase of FoxM1 expression in aged WT mice compared to vector DNA-transduced mice (**Fig. 3D**). EBA flux was drastically decreased (**Fig. 3E**) and lung MPO activity returned to a level close to basal level (**Fig. 3F**) in *FOXM1* plasmid DNA-transduced mice in contrast to vector DNA-transduced mice.

We also assessed whether the restored vascular repair and resolution of inflammation is attributable to reactivated endothelial proliferation (i.e. regeneration) in aged lungs. BrdU labeling study revealed a marked increase of EC proliferation in lungs of *FOXM1* plasmid DNA-transduced mice in sharp contrast to vector DNA-transduced mice (**Fig. 3, G** and **H**). Expression of FoxM1 target genes essential for cell cycle progression including *Cdc25c*, *Ccna2*, and *Ccnb1* was also markedly induced in lungs of *FOXM1* plasmid DNA-transduced mice (**Fig. 3I**).

To further determine if forced expression of FoxM1 in mice at very old age (e.g., 25 mos. old, equivalent to human age of ≥80 years) could still reactivate the vascular repair program, we employed our newly developed poly(lactide-co-glycolide)-*b*-poly(ethylene glycol) copolymer (PLGA-PEG)-based nanoparticles to deliver the *FOXM1* plasmid DNA. The mixture of nanoparticle:plasmid DNA was administrated retro-orbitally to 25 mos. old mice at 24h post-LPS (established injury). At 96h post-LPS, lungs were collected for EBA and MPO assays. As shown in **Fig. 3J**, lung vascular permeability measured in vector DNA-transduced mice at 96h post-LPS remained markedly elevated whereas it was greatly reduced in *FOXM1* plasmid DNA-transduced mice comparable to the observation in 19-21 mos. old mice (**Fig. 3E**). Similarly, lung MPO activity in *FOXM1* plasmid DNA-transduced mice was also markedly reduced (**Fig. 3K**), indicating normalized resolution of inflammation.

### Failure of FoxM1 Induction in pulmonary vascular ECs of elderly COVID-19 patients

To validate the potential clinical relevance of our findings in aged mice, we collected lung autopsy samples from COVID-19 patients (**Table S1**) and carried out RNAscope in situ hybridization assay to determine FOXM1 expression. FOXM1 expression in pulmonary vascular ECs was markedly induced in middle-aged COVID-19 patients but not in elderly patients (**Fig. 4**). Anti-CD31 immunostaining shows extensive disruption of the endothelial monolayer of COVID-19 patients in both middle-aged and elderly patients (**Fig. 4A**), manifesting the characteristic feature of endothelial injury of severe COVID-19 patients (*16–19*).

**Fig. 4.**
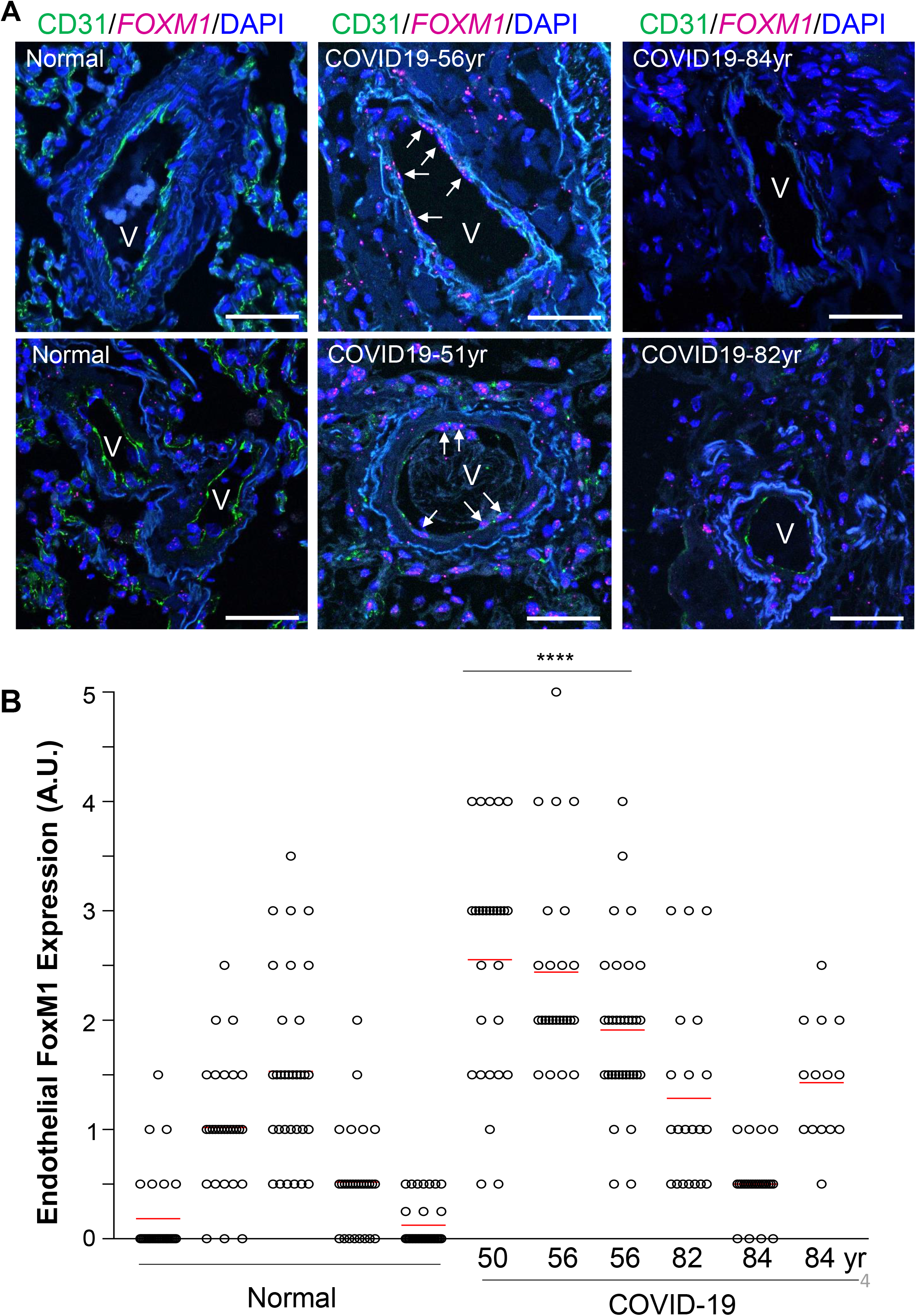
Failure of FOXM1 induction in pulmonary vascular ECs of elderly COVID-19 patients in contrast to middle-aged patients. (**A**) Representative micrographs of RNAscope in situ hybridization staining of human lung sections showing marked induction of FOXM1 expression in pulmonary vascular ECs of middle-aged COVID-19 patients but not in elderly patients. Lung autopsy tissues were collected from COVID-19 patients and healthy donors (normal) for paraffin-sectioning and immunostaining. Anti-CD31 antibody was used to immunostain ECs (green). FOXM1 mRNA expression (purple) was detected by RNAscope in situ hybridization. Nuclei were counterstained with DAPI. Arrow point to FOXM1 expressing ECs. V, vessel. Scale bar, 50 μm. (**B**) Quantification of endothelial expression of FOXM1. FOXM1 was markedly induced in ECs of middle-aged COVID-19 patients but not in elderly COVID-19 patients. FOXM1 expression was quantified in 14-33 vessels of each subject. Bars (red) represent means. **** *P* < 0.0001, Kruskal-Wallis test (non-parametric).

### Therapeutic activation of FoxM1-dependent endothelial regeneration and vascular repair in aged lungs

We next explored the possibility of pharmacological activation of FoxM1-dependent endothelial regeneration in aged lungs which will have great translational potential for treatment of ARDS and severe COVID-19 in elderly patients. Given the important role of epigenetics in aging, we focused on DNA methyl transferase inhibitor (e.g., 5-Aza 2’-deoxycytidine) and histone deacetylase inhibitor (Trichostatin A). In a preliminary study, we observed that 5-Aza 2’-deoxycytidine (i.e. Decitabine) but not Trichostatin A treatment could normalize vascular repair in aged mice following LPS challenge (data not shown). Thus, we chose to address the potential of the FDA-approved drug Decitabine. At 24h and 48h post-LPS, aged (21-22 mos. old) mice were treated with Decitabine or vehicle (PBS) and lung tissues were collected at 96h post-LPS for analyses. As shown in **Fig. 5A**, FoxM1 expression was markedly induced in lungs of Decitabine-treated mice. EBA assay demonstrated normalized vascular repair in Decitabine-treated mice in contrast to vehicle-treated mice (**Fig. 5B**). Lung MPO activity of Decitabine-treated mice was also returned to basal levels (**Fig. 5C**). However, Decitabine treatment didn’t promote vascular repair and inflammation resolution in young adult mice (**fig. S8**). Also early Decitabine treatment (e.g. at 2h post-LPS) didn’t have detrimental effects on LPS-induced lung injury (**fig. S9**), indicating a big safe window (any time after injury) for administering Decitabine to patients.

**Fig. 5.**
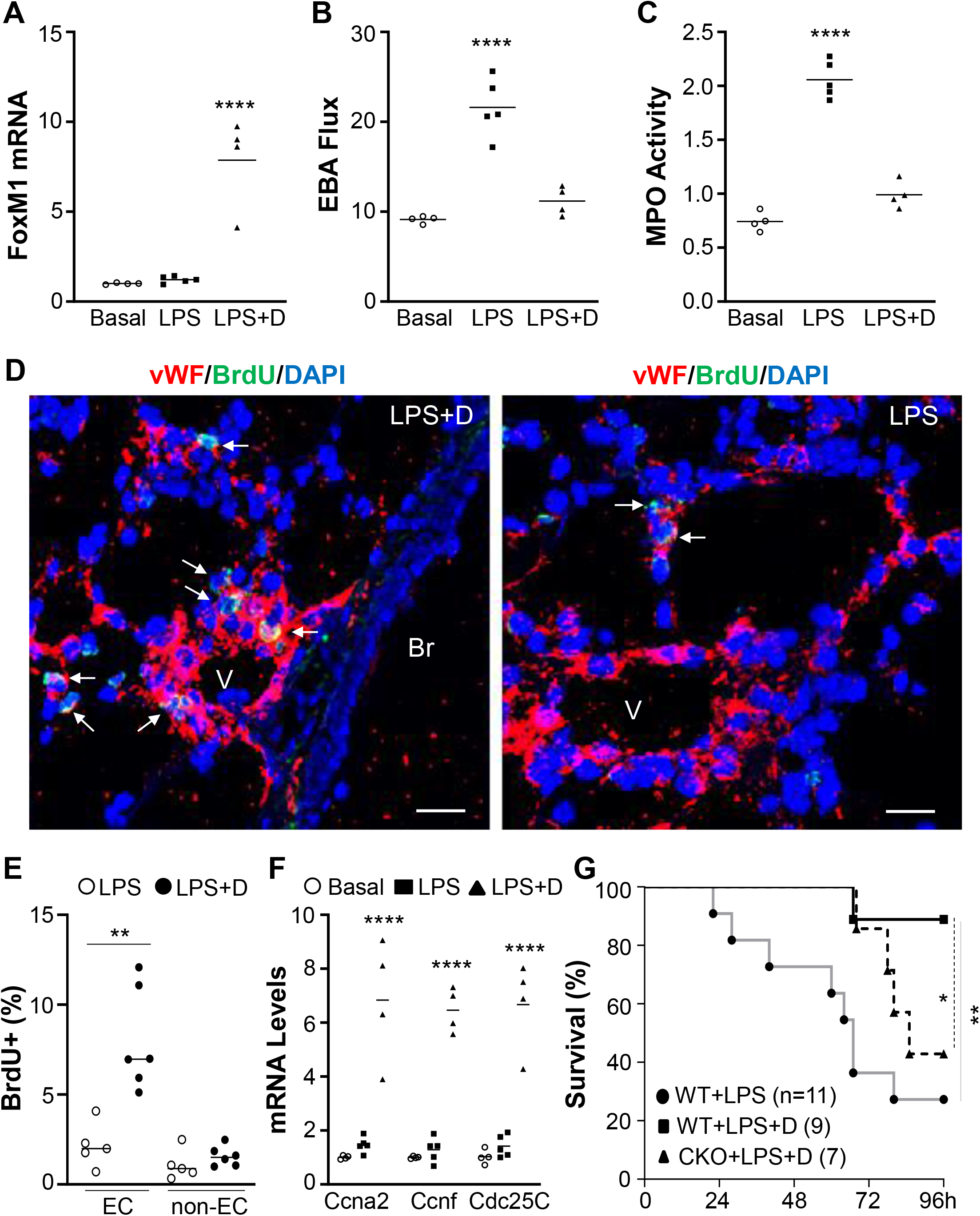
Decitabine activation of FoxM1-mediated endothelial regeneration normalized resolution of inflammatory lung injury and promoted survival of aged WT mice. Aged WT mice (21 mos.) were challenged with LPS (1.0 mg/kg, i.p.) and then treated with either Decitabine (LPS+D) or vehicle (PBS) (LPS) at 24 and 48h post-LPS (once a day). Lung tissues were collected at 96h post-LPS for various assays. (**A**) Quantitative RT-PCR analysis of FoxM1 expression. (**B**) Decitabine treatment activated lung vascular repair in aged WT mice. (**C**) Lung MPO activity of Decitabine-treated WT mice returned to levels similar to basal levels in contrast to vehicle-treated mice. (**D**) Representative micrographs of anti-BrdU staining. ECs were immunostained with anti-vWF (red). Arrows point to BrdU^+^ ECs. Br=bronchiole; V=vessel. Scale bar, 20 μm. (**E**) Quantification of BrdU^+^ ECs and non-ECs. (**F**) Quantitative RT-CPR analysis of FoxM1 target genes. (**G**) Decitabine treatment promoted survival of aged WT mice but not *Foxm1^ΔEC^* (CKO) mice. Aged WT and *Foxm1^ΔEC^* mice were challenged with LPS (0.5 mg/kg, i.p.) and then treated with Decitabine (D) or PBS at 24 and 48h post-LPS. Survival rates were recorded for 96h. \**P*<0.05, \*\**P*<0.01, \*\*\*\**P*<0.0001. One-way ANOVA (**A-C**), Student t test (**E**, **F**). Log-rank (Mantel-Cox) test (**G**).

BrdU immunostaining revealed that pulmonary vascular EC proliferation was drastically increased in Decitabine-treated aged mice, indicating reactivation of endothelial regeneration (**Fig. 5, D** and **E**). Accordingly, expression of FoxM1 target genes essential for cell cycle progression were markedly induced in lungs of Decitabine-treated aged mice (**Fig. 5F**). Decitabine treatment also markedly improved survival of aged WT mice. 80% of Decitabine-treated mice survived whereas only 20% of vehicle-treated WT mice survived at the same period (**Fig. 5G**). To determine if the survival effect was mediated by Decitabine-activated FoxM1 expression in ECs, we employed the mice with EC-specific knockout of *Foxm1* (*Foxm1^ΔEC^*) (*27*). As shown in **Fig. 5G**. Decitabine treatment had no protective effects on the survival of aged *Foxm1^ΔEC^* mice following LPS challenge.

## DISCUSSION

The present study demonstrates that in young adult mice, lung resident ECs mediate endothelial regeneration responsible for vascular repair and resulting inflammation resolution following polymicrobial sepsis-induced injury. However, aging impairs these processes leading to persistent inflammatory lung injury and high mortality in aged mice. Mechanistically, aging inhibits FoxM1-dependent endothelial regeneration and forced expression of FoxM1 by transgene or transient plasmid DNA transduction normalizes vascular repair and resolution of inflammation and promotes survival in aged mice. We also observed marked induction of FOXM1 expression in pulmonary vascular ECs of middle-aged COVID19 patients but not in elderly patients. Importantly, treatment of Decitabine activated FoxM1-dependent endothelial regeneration leading to normalized vascular repair and resolution of inflammation and enhanced survival in aged mice following sepsis. Thus, therapeutic activation of FoxM1 expression, e.g., by repurposing Decitabine may represent an effective approach for the prevention and treatment of ARDS and severe COVID-19 in elderly patients.

Studies have demonstrated that endothelial barrier dysfunction is the major contributor to lung injury and poor prognostic outcomes of sepsis, ARDS (*4, 5, 37–41*), and severe COVID19 (*16–19*). Severe inflammation such as following sepsis induces pulmonary vascular EC loss and disrupts endothelial barrier. Employing genetic lineage tracing and FACS analysis, we demonstrate that resident EC is the origin of source for lung endothelial regeneration. Our bone marrow transplantation study further excluded the contribution of bone marrow-derived cells in this regenerative process. Together, these studies provide unequivocal evidence of the exclusive contribution of lung resident ECs to endothelial regeneration in a clinically relevant sepsis model, CLP-induced polymicrobial sepsis. Employing this genetic lineage tracing mouse model, our studies have demonstrated impaired endothelial regeneration in aged lungs in contrast to young adult mice. We observed defective endothelial proliferation and vascular repair in aged lungs following lung injury induced by both polymicrobial sepsis and endotoxemia, which resulted in persistent lung vascular leaking and inflammation and thus high mortality. The impaired recovery of aged lungs following sepsis challenge is not attributable to more severe injury in aged mice than young adult mice as we employed different doses of LPS to induce similar degree of lung injury during the injury phase. We also found that mice starting at age of 12 mos. exhibited defective vascular repair and inflammation resolution which had becoming more severe with aging. These data could explain the clinical observations that the incidence and mortality of ARDS resulting from sepsis, pneumonia, and COVID-19 in elderly patients was much higher than young adult patients (*8–15*).

ECs are normally quiescent with a very low turnover rate. In response to injury, expression of some transcriptional factors is induced to activate EC proliferation (*42*). We have shown previously that FoxM1 is markedly induced in lung ECs in the recovery phase but not in the injury phase in young adult mice following LPS challenge (*27*). Here we show that FoxM1 was not induced in aged lungs following sepsis challenge, which played a causal role in aging-impaired endothelial regeneration and resolution of inflammatory injury as transgenic expression of FoxM1 prevented the defective phenotype in aged *FOXM1^Tg^* mice and promoted survival following LPS challenge. As FoxM1 was expressed ubiquitously in all cells in the *FOXM1^Tg^* mice by the *Rosa26* promoter, it is unknown if the beneficial effects were mediated by FoxM1-dependent endothelial regeneration in aged lungs. Genetic compensation of FoxM1 overexpression in *FOXM1^Tg^* mice may also contribute to the normalized vascular repair in aged mice. Thus, we designed a gene therapy-like approach to determine if transient expression of FoxM1 in ECs delivered after vascular injury could reactivate the intrinsic endothelial regeneration and vascular repair program in aged WT mice. To our surprise, transient expression of a single transcription factor in aged lungs could reactivate the intrinsic endothelial regeneration program and normalize the resolution of inflammatory lung injury in aged WT mice, even in mice at age of 25 mos., which is equivalent to human age of 80 years or older (*43*). As the plasmid DNA was transduced at 12 or 24h post-LPS challenge with established injury, this study provides unequivocal evidence of the necessary and sufficient role of restored FoxM1 expression in reactivating vascular repair and inflammation resolution in aged lungs.

Although FOXM1 expression was markedly induced in pulmonary vascular ECs of middle-aged COVID-19 patients, it was not induced in elderly COVID-19 patients. These data demonstrate the clinical relevance of our observations in aged mice. In a mouse model of SARS-CoV-2 infection, FoxM1 expression is also markedly induced in the lungs of young adult mice at 7 days after infection (*44*). Together, these studies provide strong evidence of the clinical importance of FoxM1-dependent endothelial regeneration in repairing the injured vasculature and promoting recovery and survival from severe COVID-19. Thus, identification of therapeutic agents to reactivate the aging-impaired FoxM1-dependent endothelial regeneration has great translational potential.

Our study shows that the FDA-approved drug Decitabine which is used to treat myelodysplastic syndrome can be repurposed for the prevention and treatment of ARDS and severe COVID-19 in elderly patients. Decitabine treatment of aged mice with established injury (24h post-LPS) could reactivate FoxM1 expression and endothelial proliferation in aged lungs and thus promote vascular repair and survival. Furthermore, the survival effect is mediated by endothelial FoxM1 expression as Decitabine has no survival effect on aged *Foxm1^ΔEC^* mice. However, Decitabine treatment has no effects on vascular repair in young adult mice. Our unpublished study also revealed that Decitabine had no protective effects on inflammatory lung injury during the injury phase (at 24h) following LPS challenge in young adult mice. Consistently, published studies have also shown that 5’-Aza 2’-deoxycytidine (the chemical agent of Decitabine) alone has no protective effects on LPS-induced lung vascular injury although a combination of 5’-Aza 2’-deoxycytidine and trichostatin A can inhibit LPS-induced lung vascular injury and promote survival in young adult mice, which are not ascribed to augmented vascular repair (*45, 46*). To the best of our knowledge, our data for the first time provide unequivocal evidence to support the role of Decitabine in reactivating the aging-impaired FoxM1-dependent endothelial regeneration program for vascular repair and inflammation resolution in aged mice. Intriguingly, Decitabine is under a clinical trial to test its safety and efficacy in treating severe COVID-19 patients (NCT04482621). Based on our studies, the Decitabine dose should be further optimized and the patient population should be focused on elderly patients.

In summary, we have shown aging impairs the resolution of inflammatory lung injury following sepsis challenge through inhibition of FoxM1-dependent endothelial regeneration and vascular repair. Therapeutic activation of FoxM1 in ECs could reactivate the intrinsic endothelial regeneration program for normalized resolution of inflammatory injury in aged lungs. Thus, activation of endothelial FoxM1 expression by repurposing Decitabine or nanoparticle delivery of FOXM1 plasmid DNA is a potentially effective therapeutic strategy to restore the endothelial barrier integrity and reverse lung edema in the prevention and treatment of ARDS and severe COVID-19 in elderly patients and promote survival.

## Acknowledgements

This work was supported in part by NIH grants R01HL123957, R01HL133951, R01HL140409, and R01HL148810 to Y.Y.Z.

## Author Contribution

X.H., X.Z., and Y.Y.Z. conceived the experiments. X.H., X.Z., N.M. designed, carried out experiments, and analyzed the data. X.H., X.Z., Y.Y.Z. analyzed and interpreted the data. G.M., Y.F., and D.W. provided the COVID-19 patient samples and intellectual input. X.H. wrote the manuscript. Y.Y.Z. supervised the project and revised the manuscript and is responsible for the concept.

## Supplementary Materials

### List of Supplementary Materials

Materials and Methods

Table S1

Fig. S1–S9

References (47-50)

## Materials and Methods

### Mice

*EndoSCL*-Cre^ERT2^/mTmG lineage tracing mice were generated by breeding the mice carrying a double-fluorescent reporter expressing membrane-targeted tandem dimer Tomato (mT) prior to Cre-mediated excision and membrane-targeted green fluorescent protein (mG) after excision (mTmG mice, #007676, the Jackson Laboratory) with *EndoSCL*-Cre^ERT2^ transgenic mice (C57BL/6 background) containing tamoxifen-inducible Cre-ERT2 driven by the 5’ endothelial enhancer of the stem cell leukemia locus (*47, 48*). *Foxm1* transgenic (*Foxm1^Tg^*) mice driven by the *Rosa26* promoter were described previously (*35*). *Foxm1^ΔEC^* mice with *Tie2*Cre-mediated disruption of *Foxm1* was described previously (*27*). Both male and female mice were used in the experiments. Mice at various ages (3-5 mos. old referred as young adult; 19-25 mos. old referred as aged) were used. The experiments were conducted according to NIH guidelines on the use of laboratory animals. The animal care and study protocols were approved by the Institutional Animal Care and Use Committees of Northwestern University and The University of Illinois at Chicago.

### Human specimens

All experiments with human tissue samples were performed under protocols approved by the Institutional Review Boards at the University of Chicago and Ann & Robert H. Lurie Children’s Hospital of Chicago. Postmortem lung samples were collected from diseased COVID19 patients. Control lung tissues from unused healthy donors were provided by Pulmonary Hypertension Breakthrough Initiative (PHBI) (*49*).

### Induction of lung injury

Polymicrobial sepsis was induced by cecal ligation and puncture (CLP) using a 23-gauge needle (*50*). Briefly, mice were anesthetized with inhaled isofluorane (2.5% mixed with oxygen). When the mice failed to respond to paw pinch, buprenex (0.1 mg/kg) was administered subcutaneously prior to sterilization of the skin with povidone iodine, then a midline abdominal incision was made. The cecum was exposed and ligated with a 4-0 silk tie placed 0.6 cm from the cecum tip, and the cecal wall was perforated with a 23-gauge needle. Control mice (sham) underwent anesthesia, laparotomy, and wound closure, but no cecal ligation or puncture. Following the procedure, 500 μl of prewarmed normal saline was administered subcutaneously. Within 5 min following surgery, the mice woke from anesthesia. The recovered mice subcutaneously received a second dose of buprenex at 8h post-surgery.

To induce endotoxemia, mice received a single dose of LPS (Escherichia coli 055:B5, Santa Cruz, St. Dallas, TX) by i.p. injection. The LPS dose was dependent on the age of the mice (3-9 mos. old, 2.5 mg/kg; 19-20 mos. old, 1.0 mg/kg; 21-22 mos. 0.5 mg/kg, 25 mos. old, 0.25 mg/kg). All mice were anesthetized with ketamine/xylazine (100/5 mg/kg BW, i.p.) prior to tissue collection. For the survival study, mice were treated with a single dose of LPS and monitored for 4-7 days.

### Bone marrow transplantation

Lethal irradiation was peformed with a 6MV photon beam from a linear accelerator (*27*). C57BL/6 WT mice (2 mos. old) were delivered a total of 1000 cGy at a dose rate of 100 cGy/min. At 3h following irradiation, the mice were transplanted with 10 million of bone marrow cells (200 μl of EBM2 medium) freshly isolated from *EndoSCL*-Cre^ERT2^ transgenic mice (2 mos. old) through tail vein injection. Five weeks post-transplantation, the chimeric mice were treated with tamoxifen to induce Cre-mediated GFP lableling. Three weeks later, the mice were challenged with CLP for assessment of bone marrow-derived GFP^+^ cells in lung endothelial regeneration.

### Vascular permeability assessment

The Evans blue dye-conjugated albumin (EBA) extravasation assay was performed as previously described (*50*). Briefly, mice were retro-orbitally injected with EBA at a dose of 20 mg/kg BW at 30 minutes prior to tissue collection. Lungs were perfused free of blood with PBS, blotted dry and weighed. Lung tissues were then homogenized in 1 ml PBS and incubated with 2 volumes of formamide at 60°C for 18 hours. The homogenates were centrifuged at 10, 000 × g for 30 minutes and the optical density of the supernatant was determined at 620 nm and 740 nm. The extravasated EBA in lung homogenate was presented as μg of Evans blue dye per g lung tissue.

### Myeloperoxidase assay

Following perfusion free of blood, lung tissues were collected and homogenized in 50 mmol/L phosphate buffer. Homogenates were then centrifuged at 15, 000 × g for 20 minutes at 4°C. The pellets were resuspended in phosphate buffer containing 0.5% hexadecyl trimethylammonium bromide and subjected to a cycle of freezing and thawing. Subsequently, the pellets were homogenized and the homogenates were centrifuged again. Absorbance was measured at 460 nm every 15secs for 3 minutes and data expressed as ΔOD_460_/min/g lung tissue (*27, 50*).

### FACS analysis

After perfusion free of blood with PBS, lung tissues were cut into small pieces, and incubated with 1 mg/ml collagenase A for 1 h at 37°C in a shaking water bath (200rpm). After digestion, the tissue was dispersed to a single cell preparation using the gentleMACS^TM^ Dissociator (Miltenyi Biotec, Auburn, CA) with the lung program 2. The cells were then filtered using a 40 μm Nylon cell strainer and blocked with 20% FBS for 30 min. After incubation with Fc blocker (1 μg/10^6^ cells, BD Biosciences, San Jose, CA), the cells were immunostained with anti-CD45-PB (1:800, BioLegend. San Diego, CA) and/or anti-CD31-APC (1:600, BD Biosciences) for 45 min at room temperature. Cells were then analyzed by flow cytometry (Fortessa, BD Biosciences) and sorted by flow-assisted cell sorting (Moflo Asrtios machine, Beckman Coulter, Brea, CA). mGFP- or tdTomato-labelled cells were directly analyzed with 488nm or 561nm laser wavelengths, respectively.

### Cell proliferation

At 6 h prior to tissue collection, BrdU (Sigma-Aldrich, St Louis, MO) was injected i.p. into mice at 100 mg/kg BW. Mouse lung cryosections were stained overnight with anti-BrdU (1:3, BD Biosciences, or 1:1000 Cell Signaling Technology, Danvers, MA) and incubated with Alexa Fluo 488-conjugated secondary antibody (1:200, Thermal Fisher Scientific, Waltham, MA). Lung vascular ECs were immunostained with anti-vWF (1:300, Sigma-Aldrich, St Louis, MO) and anti-CD31 (1:100, BD Bioscience) antibodies at 4°C. The sections were then incubated with Alexa Fluor 594-conjugated secondary antibodies (1:200, Thermal Fisher Scientific). The nuclei were counterstained with DAPI (Thermal Fisher Scientific). Three consecutive cryosections from each mouse lung were examined, the average number of BrdU^+^ nuclei was used.

### Molecular analysis

Total RNA was isolated using a RNeasy Mini kit including DNase I digestion (Qiagen, Valencia, CA). Following reverse transcription, quantitative RT-PCR analysis was performed using a sequence detection system (ABI ViiA 7 system; Thermal Fisher Scientific). The following primers sets were used for analysis: mouse FoxM1 primers, 5′- CACTTGGATTGAGGACCACTT-3′ and 5′ -GTCGTTTCTGCTGTGATTCC-3′; mouse cyclophilin primers, 5′ -CTTGTCCATGGCAAATGCTG-3′ and 5′- TGATCTTCTTGCTGGTCTTGC-3′. Primers for mouse *Cdc25c*, *Ccna2*, *Ccnb1*, *Tnf*, *Il6*, and *Nos2* were purchased from Qiagen. The mouse gene expression was normalized to cyclophilin.

Western blot analysis was performed using an anti-FoxM1 antibody (1:800, sc-376471, Santa Cruz Biotechnology, Santa Cruz, CA) and the same blot was incubated with anti-β-actin antibody (1:3000, BD Biosciences) as a loading control.

### RNAscope in situ hybridization assay and immunostaining

To determine FOXM1 mRNA expression in ECs of COVID-19 patient lungs and control normal donor lungs, a single-plex RNAscope in situ hybridization assay (ACD, Bio-techne, Newark, CA) combined with immunofluorescent staining for CD31 as a EC marker was carried out. Briefly, the tissue sections were baked for 1 h at 60°C, deparaffinized, and treated with H_2_O_2_ for 10 min at room temperature. Target retrieval was performed for 15 min at 100°C, followed by protease treatment for 15 min at 40°C. The sections were then hybridized with human FOXM1 probe (Cat # 446941, target region 308-1244 in NM_001243088.1, ACD, Bio-techne) for 2 h at 40°C followed by signal amplification for 30 minutes using RNAscope® Multiplex Fluorescent v2 Assay (Cat # 333110, ACD, Bio-techne) as per manufacturer’s instructions. The signal was developed by incubating the slides with TSA plus Cyanine 5 system (PerkinElmer, Waltham, MA) for 30 minutes. After RNAscope assay, the slides were incubated in blocking buffer (3% BSA, 1% FBS and 0.1% normal donkey serum) for 1 h followed by with a primary antibody against CD31 (Cat # Ab28364, Abcam, Cambridge, MA) at 4°C overnight. The sections were washed and incubated with appropriate anti-rabbit secondary antibody labeled with Alexa Fluor 488 for 1 h. The slides were then counterstained with DAPI and mounted in Prolong Gold Antifade mounting medium (ThermoFisher Scientific).

To quantify FOXM1 expression, a score system of 0-5 was used. 5 represented highest while 1 lowest expression in vascular ECs of each vessel. Fifteen 63X fields each section were randomly selected and examined.

### Imaging

Following immunostaining, lung sections were imaged with a confocal microscope system (LSM880; Carl Zeiss, Inc) equipped with a 63 × 1.2 NA objective lens. For lineage tracing studies, the cryosections were directly mounted with Prolong Gold mounting media containing DAPI.

### Transduction of plasmid DNA into vascular endothelial cells in mice

To make liposomes, a mixture comprised of dimethyldioctadecylammonium bromide and cholesterol (1:1 molar ratio) was dried using a Rotavaporator (Brinkmann), and dissolved in 5% glucose followed by 20 min sonication as described previously (*36, 50*). The complex consisting of plasmid DNA expressing human *FOXM1* under the control of human *CDH5* promoter or empty vector and liposomes was combined at a ratio of 1 μg of DNA to 8 nmol of liposomes. The DNA/liposome complex (50 μg of DNA/mouse) was injected into the retro-orbital venous plexus at 12h post-LPS challenge.

In a separate study, mixture of nanoparticle:plasmid DNA (15 μg DNA/mouse) was administered retro-orbitally to mice of 25 mos. old at 24h post-LPS. The novel nanoparticle (patent pending) was poly(lactide-co-glycolide) (PLGA)-based with highly efficient gene delivery to vascular ECs after retro-orbital administration.

### Statistical analysis

Data distribution normality was first assessed by Shapiro-Wilk test. Statistical significance was determined by one-way *ANOVA* with a Dunnett post hoc analysis that calculates P values corrected for multiple comparisons for equal variance or Kruskal-Wallis test for unequal variance using Prism 7 (Graphpad Software, Inc. La Jolla, CA). Two-group comparisons were analyzed by the unpaired 2-tailed *t* test for equal variance or by Mann-Whitney test for unequal variance. Statistical analysis of the survival study was performed with the log-rank (Mantel-Cox) test. *P* < 0.05 denoted the presence of a statistically significant difference. All bars in dot plot figures represent means.

---

**Table S1.** Patient demographic information.

|  | Age<br>Yr | Sex | Hospital<br>days | ICU<br>days | Respiratory<br>symptom |
| --- | --- | --- | --- | --- | --- |
| Normal 1 | 57 | M | N/A | N/A | N/A |
| Normal 2 | 43 | F | N/A | N/A | N/A |
| Normal 3 | 60 | F | N/A | N/A | N/A |
| Normal 4 | 54 | M | N/A | N/A | N/A |
| Normal 5 | 34 | F | N/A | N/A | N/A |
| COVID-19-1 | 51 | F | 12 | 12 | yes |
| COVID-19-2 | 56 | M | 13 | 13 | yes |
| COVID-19-3 | 56 | F | 41 | 41 | No |
| COVID-19-5 | 82 | F | 13 | 13 | yes |
| COVID-19-6 | 84 | M | 1 | 0.5 | yes |
| COVID-19-7 | 84 | F | 4 | 0 | no |

## Supplemental Figures and Figure Legends

**fig. S1.**
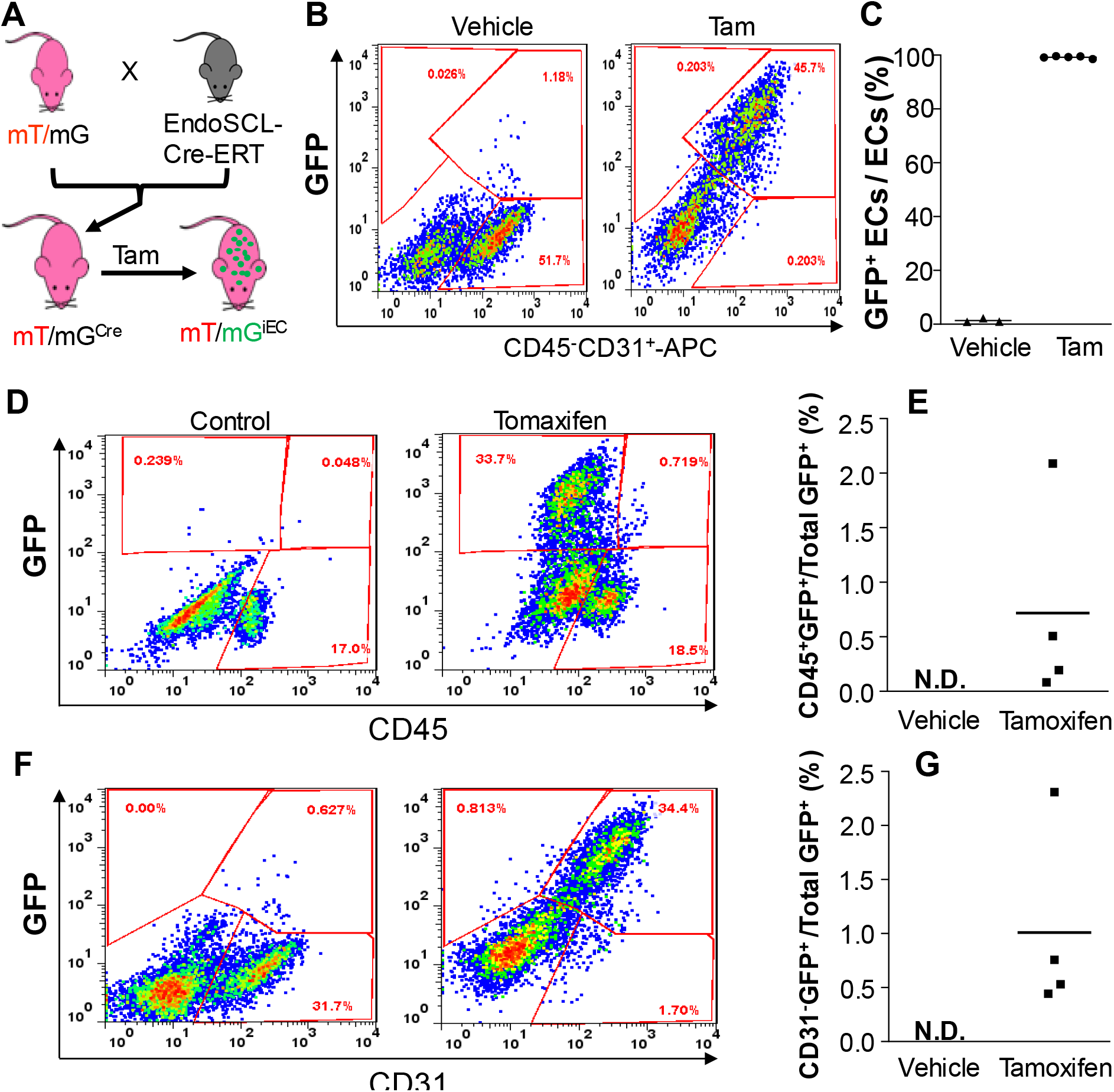
Flow cytometry analysis demonstrating highly efficient EC-specific labeling with few non-ECs are GFP^+^. (**A**) Schematic illustration of the lineage-tracing strategy. Tam=Tamoxifen. (**B**) Flow cytometry analysis of GFP^+^ cells and ECs (CD45^−^CD31^+^) in lungs of young adult mice. (**C**) Quantification of lung GFP^+^ ECs in young adult *EndoSCL*-Cre^ERT2^/mTmG mice demonstrating 95% of labeling efficiency. At 1 mo. post-tamoxifen or vehicle treatment, lung tissues were collected for isolation of cells which were then immunostained with anti-CD45 and anti-CD31 antibodies. CD45^−^ cells were gated for CD31^+^ and GFP^+^ analysis. (**D**, **E**) Few CD45^+^ cells (leukocytes) are GFP^+^ in *EndoSCL*-Cre^ERT2^/mTmG mice after tamoxifen treatment. (**F**, **G**) Few CD31^−^ cells (non-ECs) are GFP^+^ in *EndoSCL*-Cre^ERT2^/mTmG mice after tamoxifen treatment. Bars represent means. N.D., not detected.

**fig. S2.**
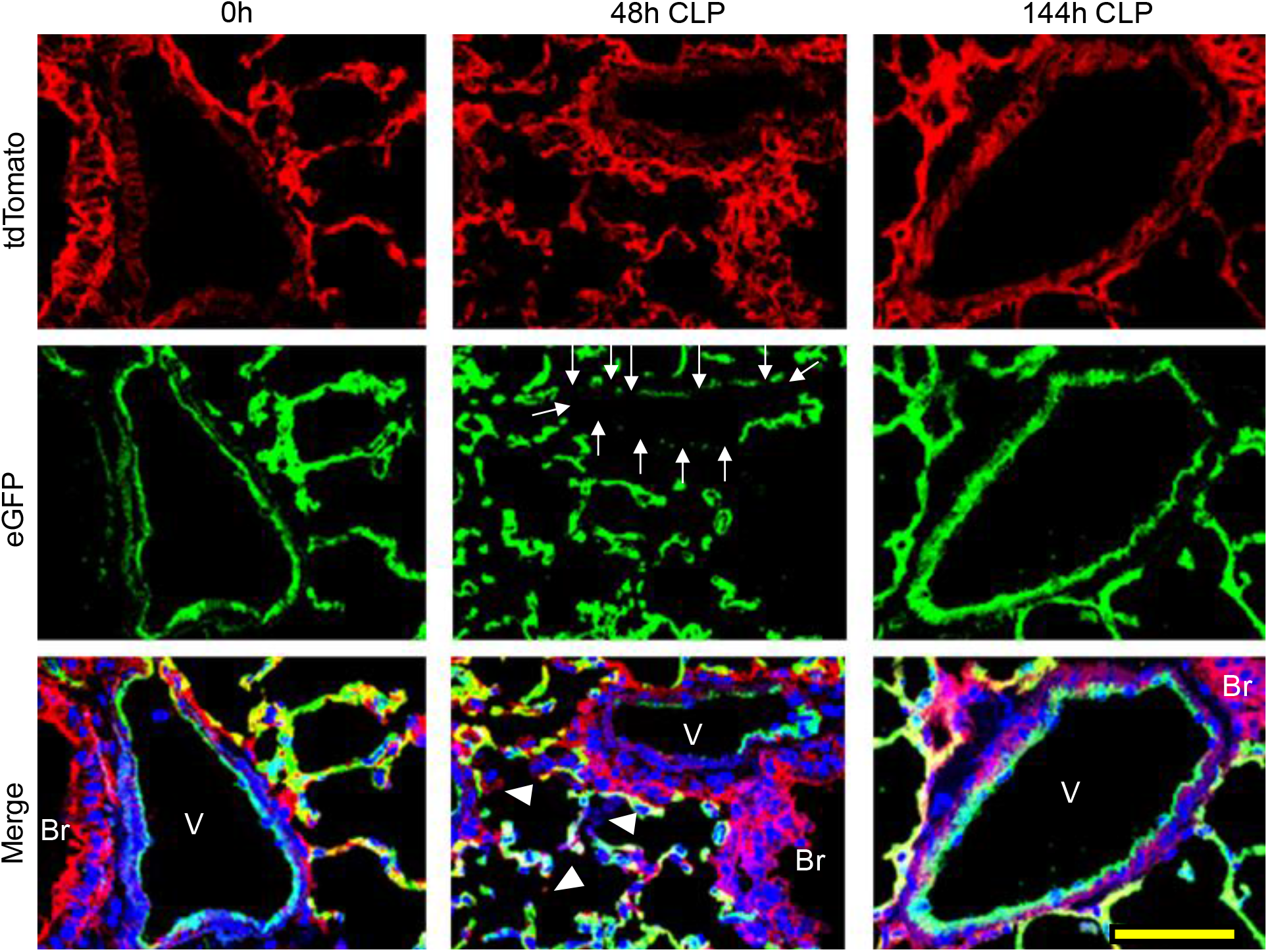
Representative confocal images of lungs of young adult mice showing changes in GFP-labeled ECs. At 48h post-CLP, loss of GFP^+^ ECs was evident in pulmonary vessel (arrows). At 144h post-CLP, vascular integrity was fully recovered evident by intact green lining as seen in Sham control lungs. Red, tdTomato^+^ (non-ECs); Green, GFP^+^ cells (ECs). Blue, DAPI (nuclei). Br, bronchiole; V, vessel. Scale bar, 20 μm.

**fig. S3.**
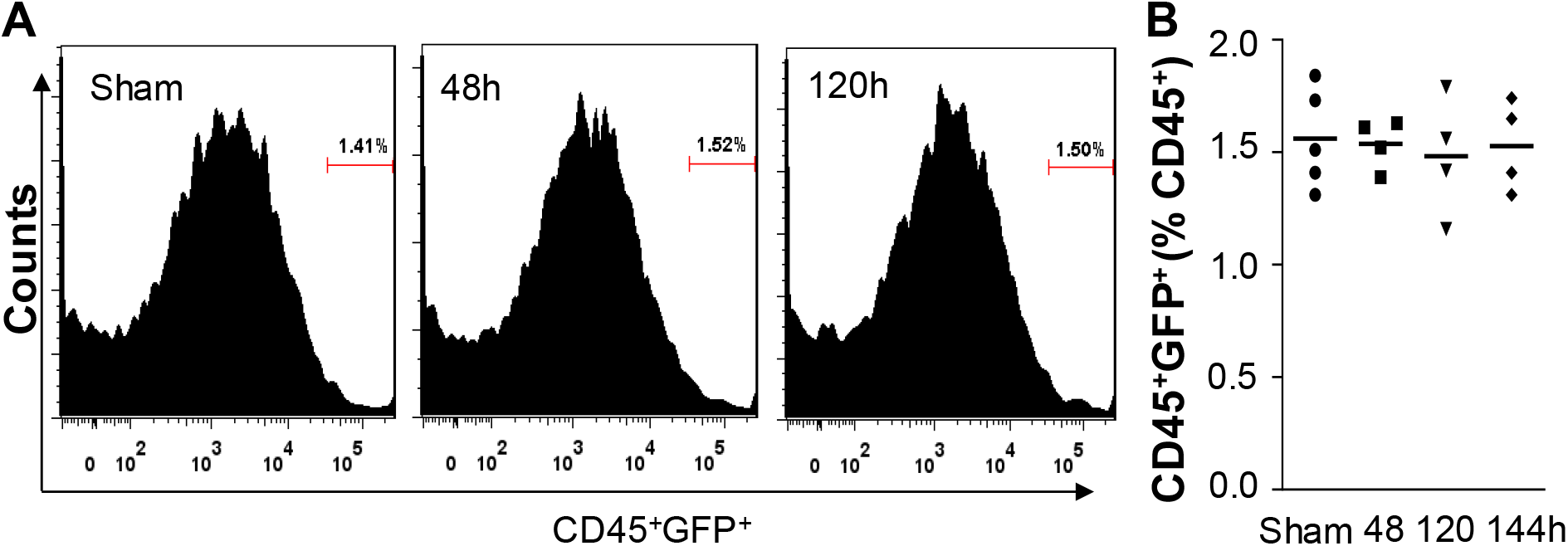
Flow cytometry analysis demonstrating CD45^+^/GFP^+^ cells were not involved in endothelial regeneration following polymicrobial sepsis. At various times following CLP challenge, lungs from tamoxifen-treated *EndoSCL*-Cre^ERT2^/mTmG mice were collected for cell isolation and the cells were immunostained with anti-CD45 antibody for flow cytometry analysis. Sham, 144h post-Sham. Bars represent means.

**fig. S4.**
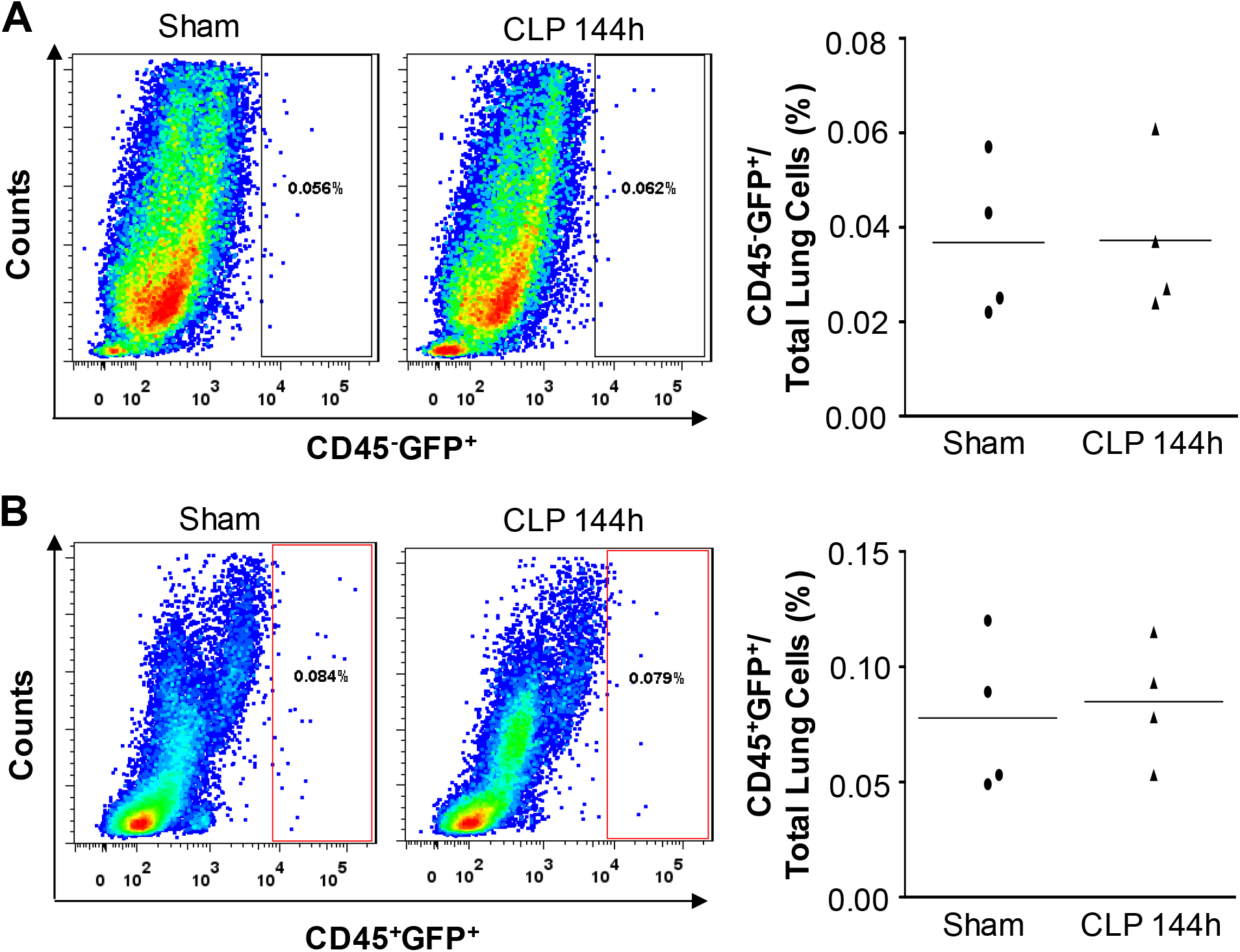
Bone marrow transplantation demonstrating little contribution of bone marrow-derived cells in lung endothelial regeneration. Bone marrow cells isolated from mTmG/*EndoSCL*-Cre^ERT2^ mice were transplanted to lethally irradiated C57BL/6 WT mice (2 mos. old) to generate chimeric mice. Upon tamoxifen treatment, bone marrow-derived *EndoSCL-Cre^+^* cells were labeled with GFP in these chimeric mice. FACS analysis shows that the percentages of CD45^−^GFP^+^ cells (e.g. ECs) and CD45^+^GFP^+^ cells in lungs of the chimeric mice at 144h post-CLP challenge were similar to that of Sham, demonstrating bone marrow-derived GFP+ cells didn’t contribute to endothelial regeneration. Bars represent means.

**fig. S5.**
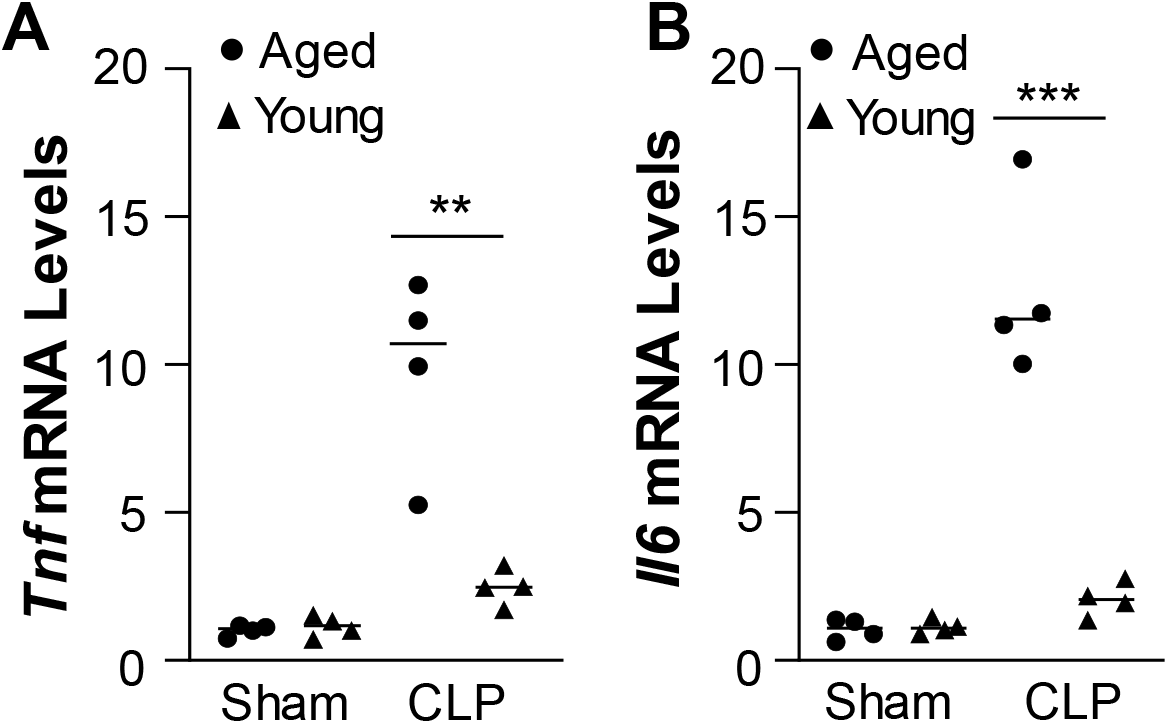
Quantitative RT-PCR analysis showing marked increases of expression of proinflammatory genes in lungs of aged mice at 96h post-CLP compared to young mice. Bars represent means. ** *P* < 0.01. Unpaired *t* test.

**fig. S6.**
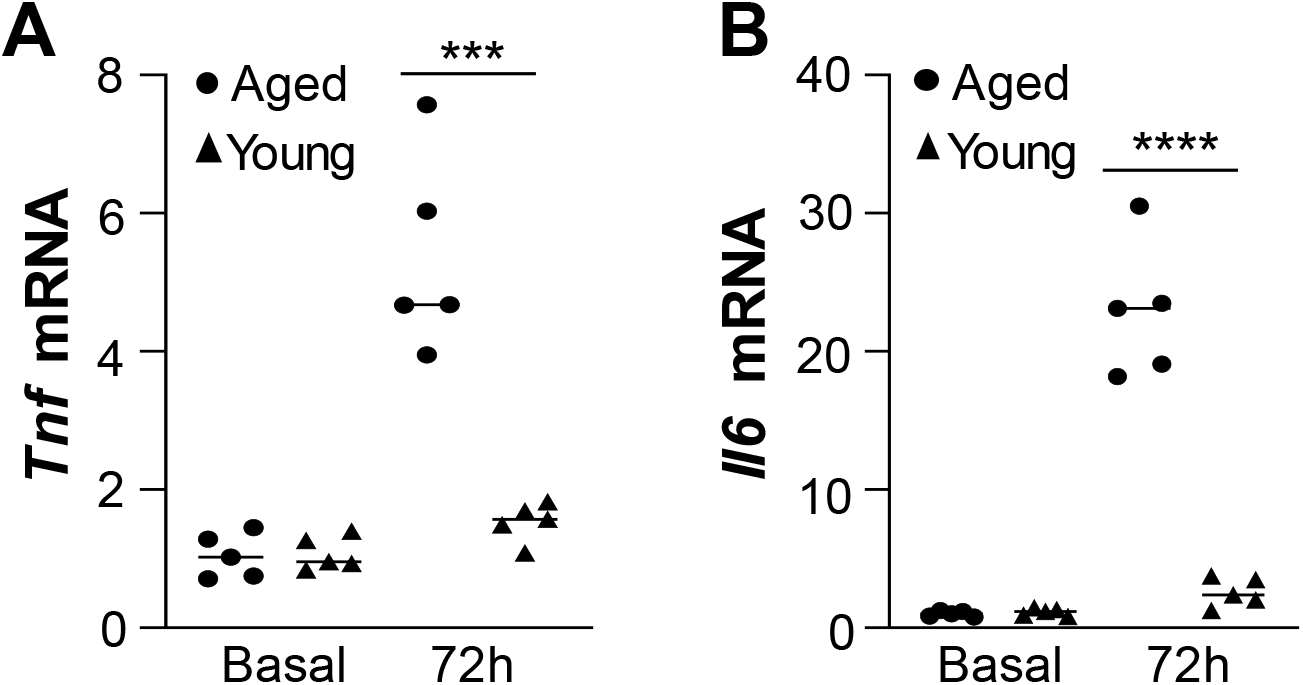
Quantitative RT-PCR analysis of expression of proinflammatory genes in mouse lungs. At 72h post-LPS, lung tissues were collected for RNA isolation followed by quantitative RT-CPR analysis. Bars represent means. \*\*\**P*<0.001; \*\*\*\**P*< 0.0001. Unpaired *t* test.

**fig. S7.**
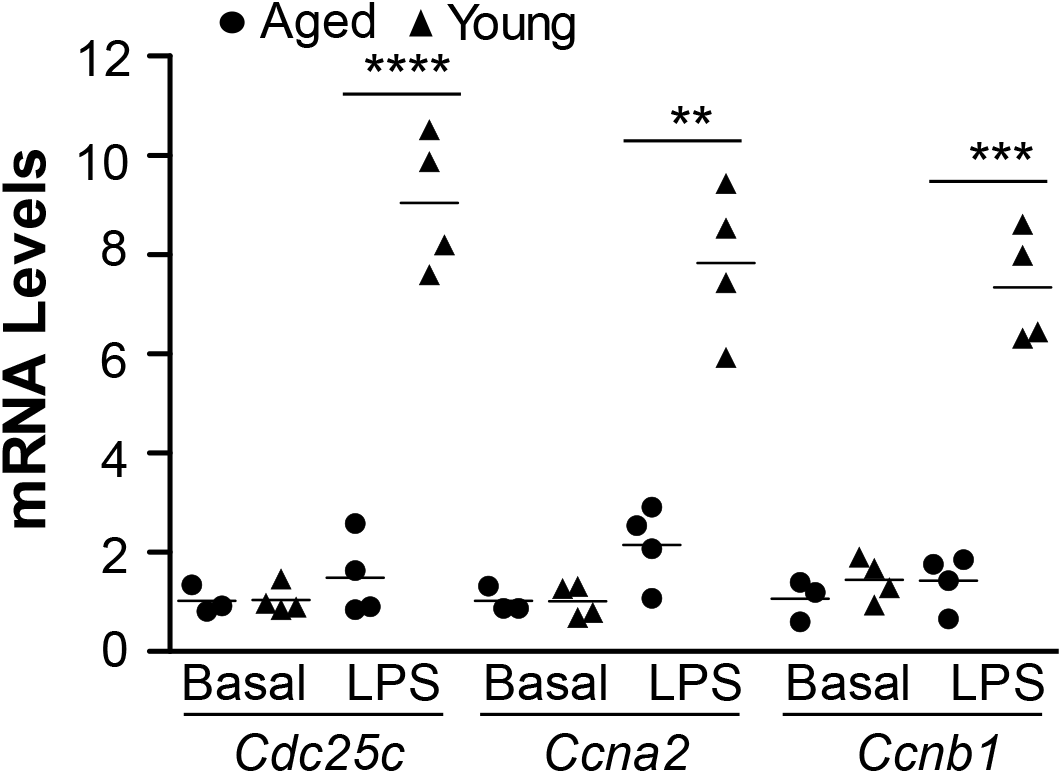
Quantitative RT-PCR analysis of expression of FoxM1 target genes essential for cell cycle progression. At 72h post-LPS, lung tissues were collected for RNA isolation. Expression of FoxM1 target genes essential for cell cycle progression was determined by quantitative RT-PCR analysis. Bars represent means. \*\**P*<0.01; \*\*\**P*<0.001. Unpaired *t* test.

**fig. S8.**
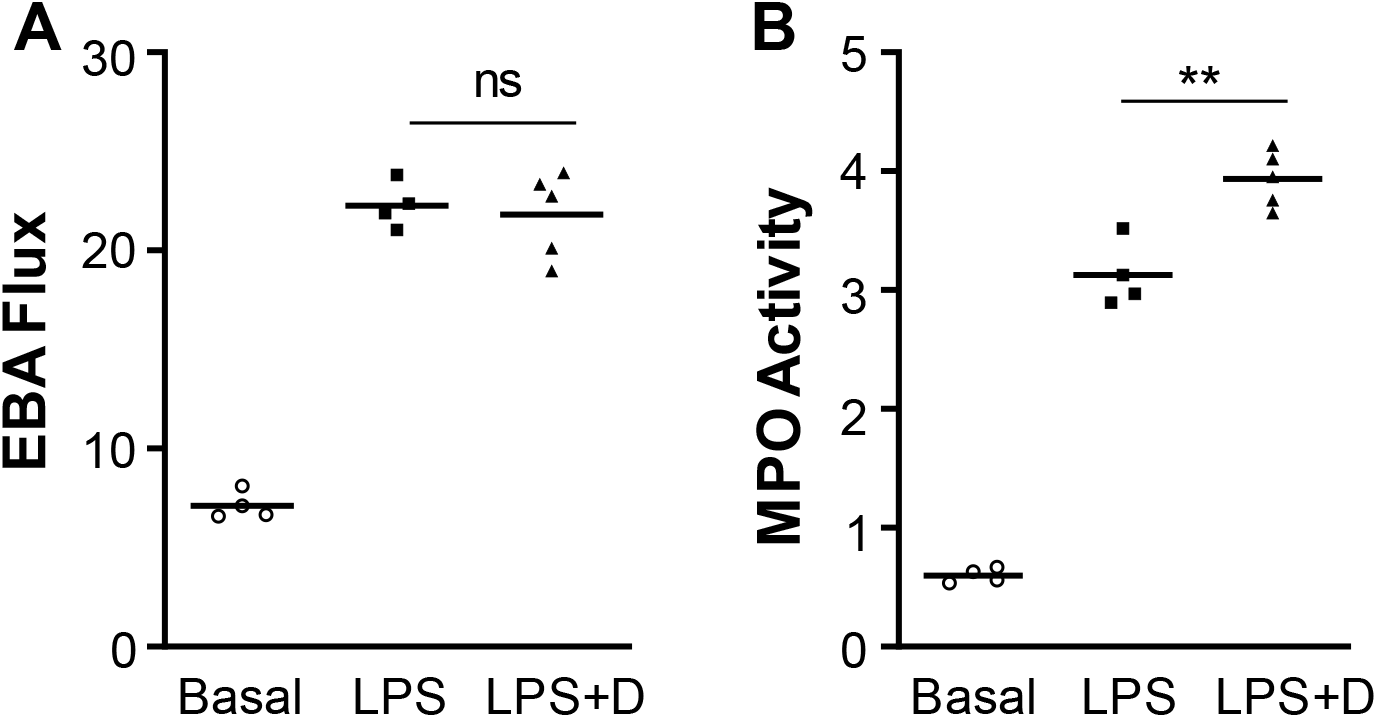
Decitabine treatment didn’t promote vascular repair in young adult mice. Mice at age of 3-4 mos. were challenged with LPS (2.5 mg/kg, i.p.) then treated with decitabine (LPS+D) or vehicle (PBS) at 4 h and 24 h post-LPS. Lung tissues were collected at 52 h for EBA flux (**A**) and MPO activity (**B**) assessment. Bars represent means. ** *P* < 0.01. Unpaired t test. ns, not significant.

**fig. S9.**
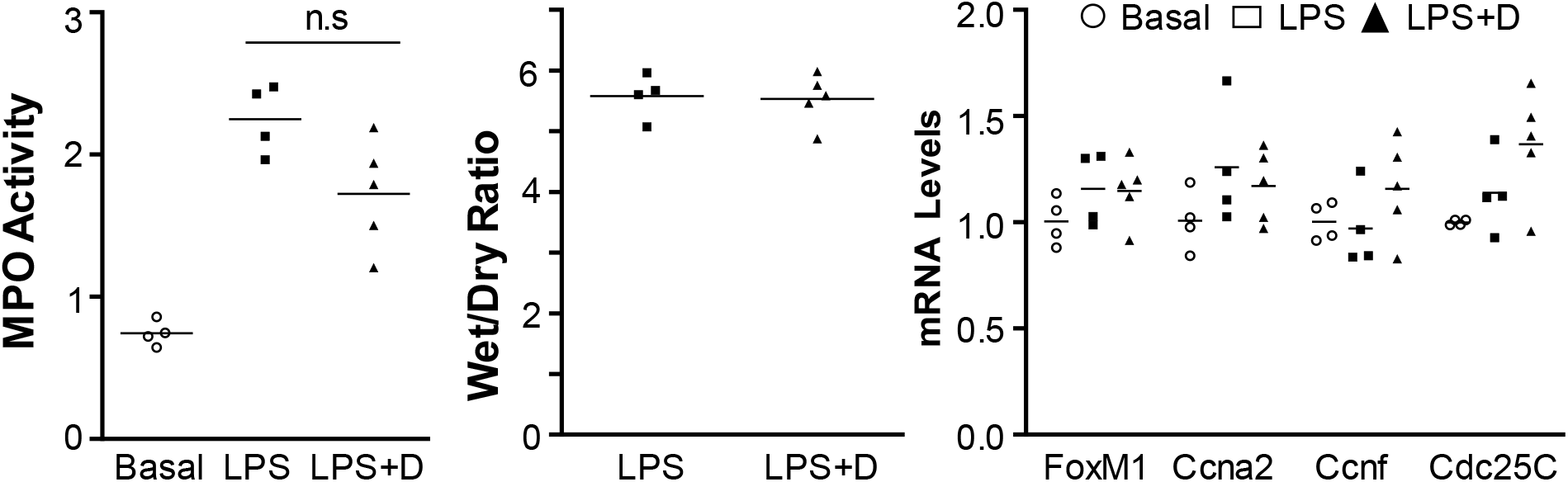
Early treatment of Decitabine didn’t induce detrimental effects on LPS-induced inflammatory lung injury in aged WT mice. Aged WT mice (21 mos. old) were challenged with LPS and then treated with either Decitabine (LPS+D) or PBS (LPS) at 2h post-LPS. Lung tissues were collected at 8h post-LPS for MPO activity assay (**A**), weight/dry weight ratio (**B**), and gene expression (**C**). Expression of FoxM1 and its target genes was not induced at this injury phase. Bars represent means. ns, not significant.

